# Large hydrogen isotope fractionations distinguish nitrogenase-derived methane from other sources

**DOI:** 10.1101/2020.04.10.036657

**Authors:** Katja E. Luxem, William D. Leavitt, Xinning Zhang

**Author notes:** **Corresponding Authors:** Katja Luxem, Xinning Zhang, Department of Geosciences (M47 Guyot Hall), Princeton University, Princeton NJ 08540.

## Abstract

Nitrogenase is the main source of natural fixed nitrogen for the biosphere. Two forms of this metalloenzyme, the vanadium (V) and iron (Fe)-only nitrogenases, were recently found to reduce small amounts of carbon dioxide into the potent greenhouse gas methane. Here we report carbon and hydrogen stable isotopic compositions and fractionations of methane generated by V- and Fe-only nitrogenases in the metabolically versatile nitrogen fixer *Rhodopseudomonas palustris*. The stable carbon isotope fractionation imparted by both forms of alternative nitrogenase are within the range observed for hydrogenotrophic methanogenesis (^13^α_CO2/CH4_ = 1.051 ± 0.002 for V-nitrogenase and 1.055 ± 0.001 for Fe-only nitrogenase, mean ± SE). In contrast, the hydrogen isotope fractionations (^2^α_H2O/CH4_ = 2.071 ± 0.014 for V-nitrogenase and 2.078 ± 0.018 for Fe-only nitrogenase) are the largest of any known biogenic or geogenic pathway. The large ^2^α_H2O/CH4_ shows that the reaction pathway nitrogenases use to form methane strongly discriminates against ^2^H, and that ^2^α_H2O/CH4_ distinguishes nitrogenase-derived methane from all other known biotic and abiotic sources. These findings on nitrogenase-derived methane will help constrain carbon and nitrogen flows in microbial communities and the role of the alternative nitrogenases in global biogeochemical cycles.

**Importance:** All forms of life require nitrogen for growth. Many different kinds of microbes living in diverse environments make inert nitrogen gas from the atmosphere bioavailable using a special protein, *nitrogenase*. Nitrogenase has a wide substrate range, and in addition to producing bioavailable nitrogen, some forms of nitrogenase also produce small amounts of the greenhouse gas methane. This is different from other microbes that produce methane to generate energy. Until now, there was no good way to determine when microbes with nitrogenases are making methane in nature. Here, we developed an isotopic fingerprint that allows scientists to distinguish methane from microbes making it for energy versus those making it as a byproduct of nitrogen acquisition. With this new fingerprint, it will be possible to improve our understanding of the relationship between methane production and nitrogen acquisition in nature.

## Introduction

Microorganisms produce over half of global methane emissions (1). Fermentative and hydrogenotrophic methanogens are the most significant microbial producers of this potent greenhouse gas (1, 2). Their metabolic pathways occur exclusively within anaerobic Archaea and involve multiple enzymes working together in series, including the obligatory methyl-coenzyme M reductase (*mcr*) enzyme. Its primary function is for catabolism, with methane production thought to occur only after other more favorable electron acceptors, like oxygen, nitrate, or sulfate have been consumed (3–5). Over the past decade, it has been recognized that minor additional contributions of methane derive from the demethylation of organophosphonates (c.f. (6–8)) and from inefficient Wood-Ljungdahl pathway carbon fixation (9). Most recently, it was discovered that some forms of the metalloenzyme nitrogenase also reduce carbon dioxide straight into methane (10). Nitrogenases are the only biological source of newly fixed nitrogen to the biosphere, and prior to industrial reduction of dinitrogen, were the primary source of nitrogen to life on Earth (11, 12). The discovery of biological methane production by nitrogenase expands the known range of organisms and environments in which methane production is possible.

Nitrogenase is known primarily for its ability to reduce inert dinitrogen (N_2_) gas into ammonia, a process known as nitrogen fixation. This biological nitrogen source plays a critical role in ecosystem fertility. Nitrogenase is generally considered a promiscuous enzyme because it can reduce a variety of carbon containing compounds in addition to N_2_ (13–17). For example, the iron (Fe)-only nitrogenase isoform can convert carbon monoxide into hydrocarbon chains, a reaction which may have been important for early forms of life (15). In addition, all forms of nitrogenase reduce acetylene to ethylene (18–21), which is the basis for the most commonly used acetylene reduction method to measure nitrogen fixation rates in the laboratory and field (22–24). The recent discovery that some forms of nitrogenase can reduce carbon dioxide to methane (10) is significant because, unlike acetylene and carbon monoxide, carbon dioxide is ubiquitous in nature.

The vanadium (V-) and Fe-only nitrogenase isoforms, which were shown to produce the most byproduct methane of the various nitrogen isoforms (10), are found in both the bacterial and archaeal domains and are widespread in nature (25–30). In addition, certain artificial mutations near the active site of the molybdenum (Mo)-nitrogenase enabled this more common isoform to produce methane (31, 32). These findings beg the question of whether and to what extent nitrogenase is an important methane source in certain environments, and how to distinguish nitrogenase-derived methane from other sources. Previous research has established that each form of nitrogenase imparts a characteristic nitrogen or carbon isotope fractionation during N_2_ (33) or acetylene (26) reduction, respectively. The stable isotopes of carbon and hydrogen are commonly used to differentiate (‘fingerprint’) different sources of methane (2, 8, 34–39). To determine what characteristic carbon and hydrogen isotope fractionations are associated with methane production by the different nitrogenases, we cultivated V- and Fe-nitrogenase utilizing strains of the anoxygenic photoheterotroph *Rhodopseudomonas palustris* under nitrogen-fixing conditions. We find that the carbon isotope fractionations are large, yet similar to those of canonical anaerobic methanogens. Conversely, the hydrogen isotope fractionation values are the largest of any methane production pathway on record. This unique hydrogen isotopic fingerprint allows us to differentiate nitrogenase-derived methane from other sources, and provides insight into the mechanism of proton delivery to nitrogenase.

## Results and Discussion

### Isotope fractionation by nitrogenase during methane production

Different methane sources are commonly associated with characteristic stable isotope fractionations that can help distinguish between different biogenic, geogenic and thermogenic sources (2, 36, 39). To determine the stable isotopes associated with methane production by nitrogenase, we grew mutant strains of the anoxygenic photoheterotroph *Rhodopseudomonas palustris* CGA009 that exclusively utilize either the Mo-nitrogenase, V-nitrogenase or Fe-only nitrogenase for nitrogen fixation (10, 40, 41). The Mo-nitrogenase strain did not produce detectable methane during batch culture incubation through stationary phase in Balch tubes (data not shown). The V- and Fe-only nitrogenase strains both produced methane, with the Fe-only nitrogenase strain producing over an order of magnitude more methane than the V-nitrogenase strain (Fig. 1). For the Fe-nitrogenase strain, methane production per cell was higher later during growth. We measured the carbon and hydrogen isotopic compositions of methane and fractionations relative to carbon dioxide (CO_2_/CH_4_) and water (H_2_O/CH_4_), as produced by the V- and Fe-only nitrogenases across a range of cell densities (OD_660_ ~ 0.3 to 1.3, from early log to stationary phase), temperatures (14 to 30°C), carbon substrates (succinate and acetate), and growth medium pH (from 6.2 to 6.8 at inoculation).

**Fig. 1.**
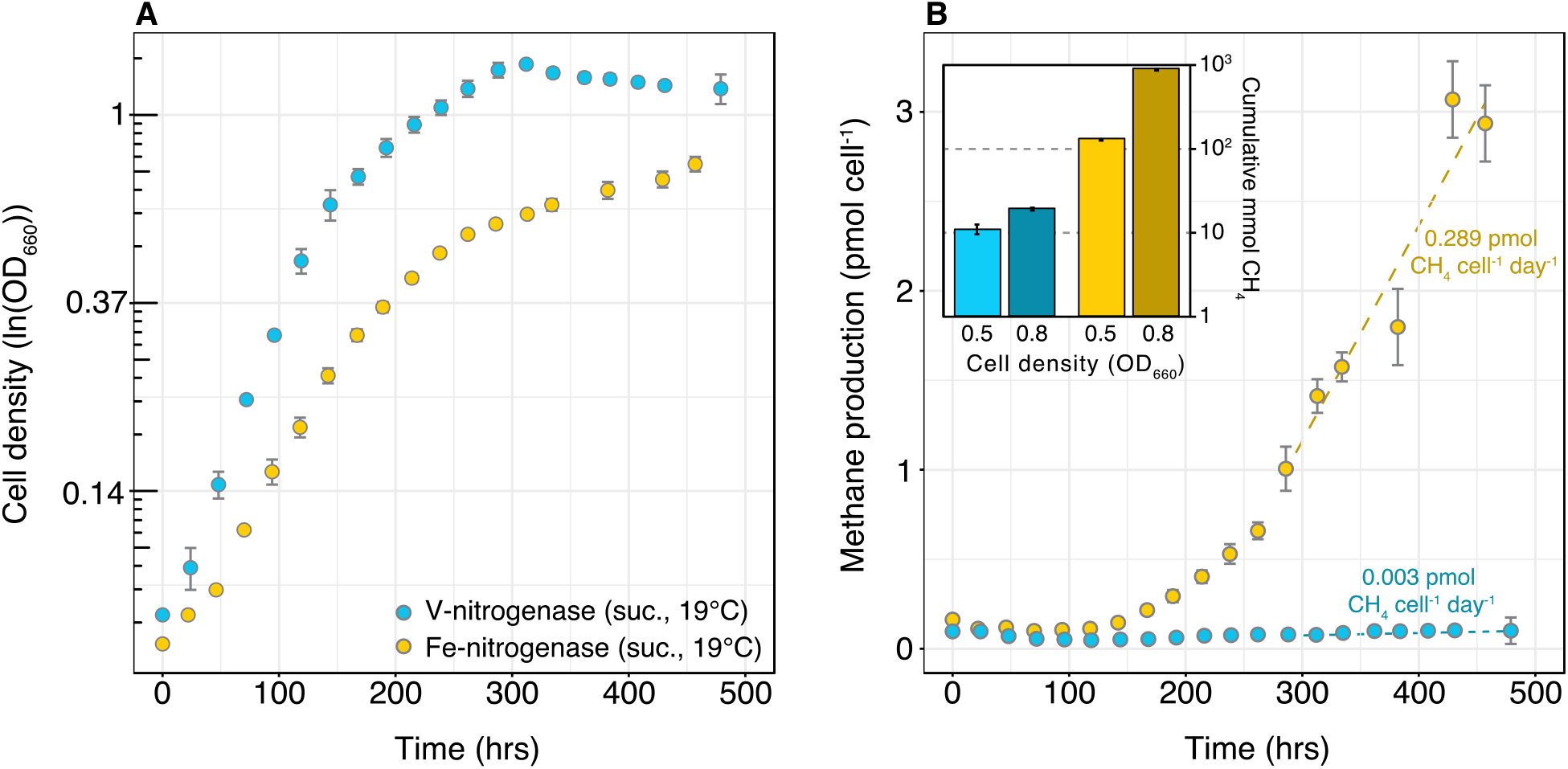
Growth dynamics and methane yields of nitrogenase strains. During growth (**A**) on succinate at 19°C, the *R. palustris* V- and Fe-only nitrogenase strains produced methane (**B)**. The Fe-only nitrogenase strain produced >10-fold more methane in the headspace than the V-nitrogenase strain. For the Fe-nitrogenase strain, methane production per cell is greater at higher cell densities. Error bars show the standard error of three biological replicates. Dissolved methane is not included.

We discovered that methane produced by the V- and Fe-only nitrogenases is highly depleted in deuterium relative to other natural sources (Fig. 2). Growth on medium with substrate water δ^2^H of ~ –40₀ yielded methane with δ^2^H values ranging from −473 to −560%₀. To our knowledge, this is the most deuterium-depleted hydrogen isotope ratio measured for natural methane sources to date. The methane carbon isotopic composition, which varied from δ^13^C = −73.0 to −97.1%₀ for substrate CO_2_ of ~ −30%₀, falls within the range observed for hydrogenotrophic methanogenesis (2) but is distinct from other abiogenic (36) and non-traditional biotic sources (8).

**Fig. 2.**
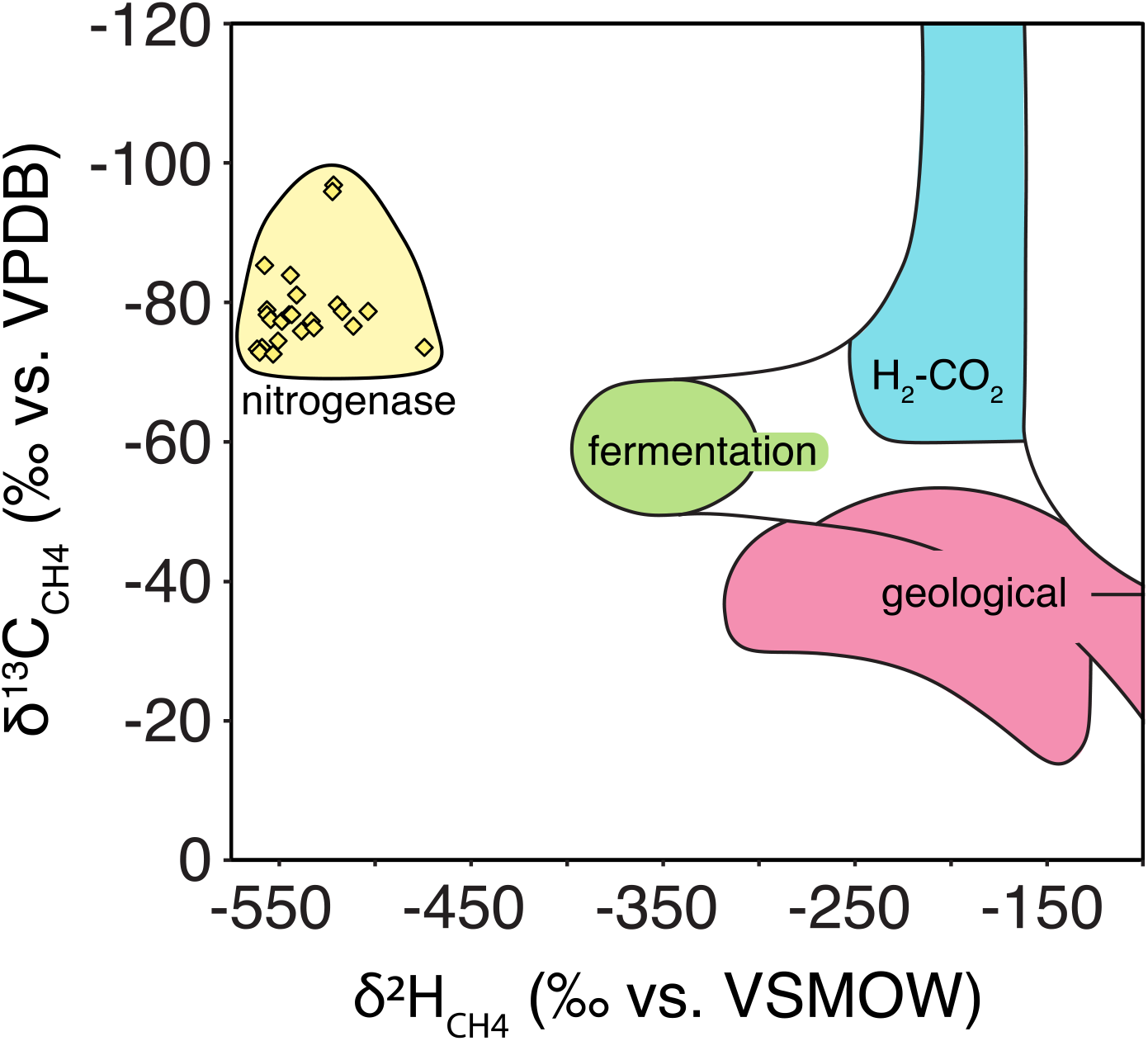
Nitrogenase-derived methane has a unique stable isotopic composition. The stable isotopic composition of methane produced by nitrogenase (yellow) can be distinguished from other natural methane sources due to its more depleted hydrogen isotopic composition. Individual datapoints from this study are shown as diamonds (♢, n = 31). The observed range for fermentative (green), hydrogenotrophic (blue) and geological (red) methane sources were taken from (90), though we note that these boundaries are not absolute (e.g. (36)).

Attributing methane isotope ratios to specific pathways becomes more reliable when the isotopic composition of source water and carbon are also considered (34, 36). In our experiments, manipulation of growth medium δ^2^H over a ~ 600%₀ range, from −30 to 550%₀, resulted in a constant, statistically indistinguishable fractionation of ^2^α_H2O/CH4_ = (δ^2^H_H2O_+1000)/(δ^2^H_CH4_+1000) = 2.047 ± 0.016 calculated for individual samples, ^2^α_H2O/CH4_ = 2.056 ± 0.057 calculated using the slope, and ^2^α_H2O/CH4_ = 2.050 ± 0.019 calculated using the intercept (*p* = 0.9; Fig. 3). The hydrogen isotope fractionations (1.820 ≤ ^2^α_H2O/CH4_ ≤ 2.199) measured for methane production by V- and Fe-only nitrogenase over a range of temperatures and growth conditions are substantially higher than the largest fractionations observed for traditional microbial methanogenesis pathways, which are around ^2^α_H2O/CH4_ ~1.45 for acetoclastic (42) and hydrogenotrophic (34) methanogenesis (Figs. 4A, 5). In fact, depending on the substrate concentrations and environmental conditions like temperature, the hydrogen isotope fractionation for these traditional methane-forming pathways is often even lower than ^2^α_H2O/CH4_ =1.45 (34, 42, 43). Our data indicate that a large hydrogen isotope fractionation of ^2^α_H2O/CH4_ ~ 2.070 is characteristic of methane production by nitrogenase and distinguishes methane produced by nitrogenase from other biogenic and abiogenic pathways.

**Fig. 3.**
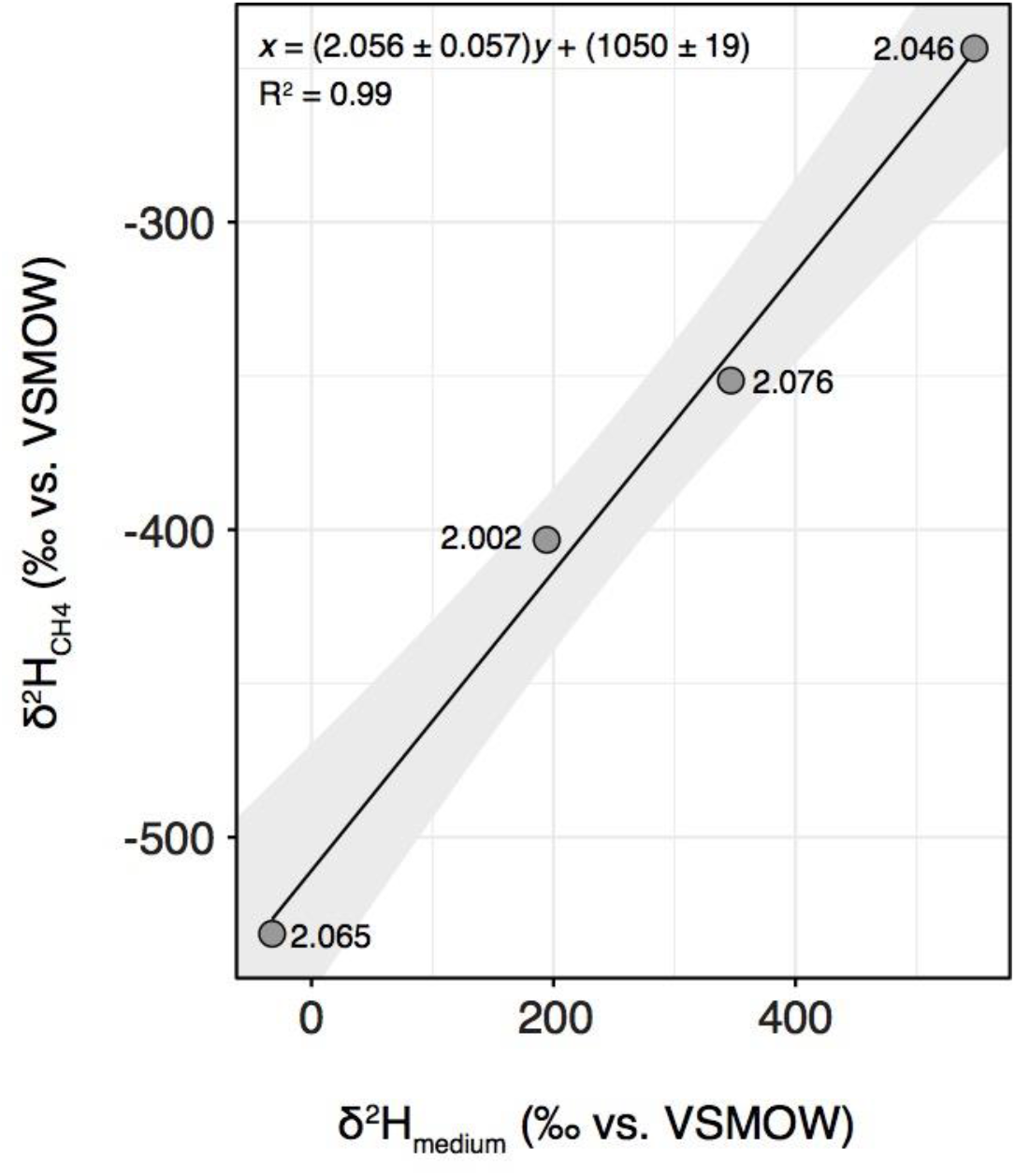
Hydrogen isotope fractionation does not depend on water isotopic composition. Regression of δ^2^H values for source water versus methane show that hydrogen isotope fractionation (^2^α_H2O/CH4_) is constant over a 600%₀ range for the Fe-only nitrogenase strain grown at 19°C on succinate. The hydrogen isotope fractionation calculated using the slope (^2^α_H2O/CH4_ = 2.056 ± 0.057, mean ± SE), intercept (^2^α_H2O/CH4_ = 1050/1000 + 1 ± 19/1000 = 2.050 ± 0.019) and individual samples (^2^α_H2O/CH4_ = 2.047 ± 0.016) is indistinguishable (*p* ≥ 0.9). The values next to each data point are the calculated fractionations for individual samples and the shaded area shows the 95% confidence interval for the regression. N.B. The convention used for individual samples is for substrate over product, whereas the regression line was calculated as product over substrate. The regression calculated for substrate over product, *x* = (2.043 ± 0.117) *y* + (1045 ± 46), is statistically indistinguishable (*p* ≥ 0.9).

Like the carbon isotopic composition, the carbon isotope fractionation measured for nitrogenase (1.045 ≤ ^13^α_CO2/CH4_ = (δ^13^CC_O2_+1000)/(δ^13^CC_H4_+1000) ≤ 1.062) falls within the range observed for hydrogenotrophic methanogenesis (1.030 ≤ ^13^α_CO2/CH4_ ≤ 1.080; (37); Fig. 4B). Mechanistically, it is possible that the similarity in carbon isotope fractionation between these two pathways is due to the similarity in substrate (CO_2_) and electron requirements.

**Fig. 4.**
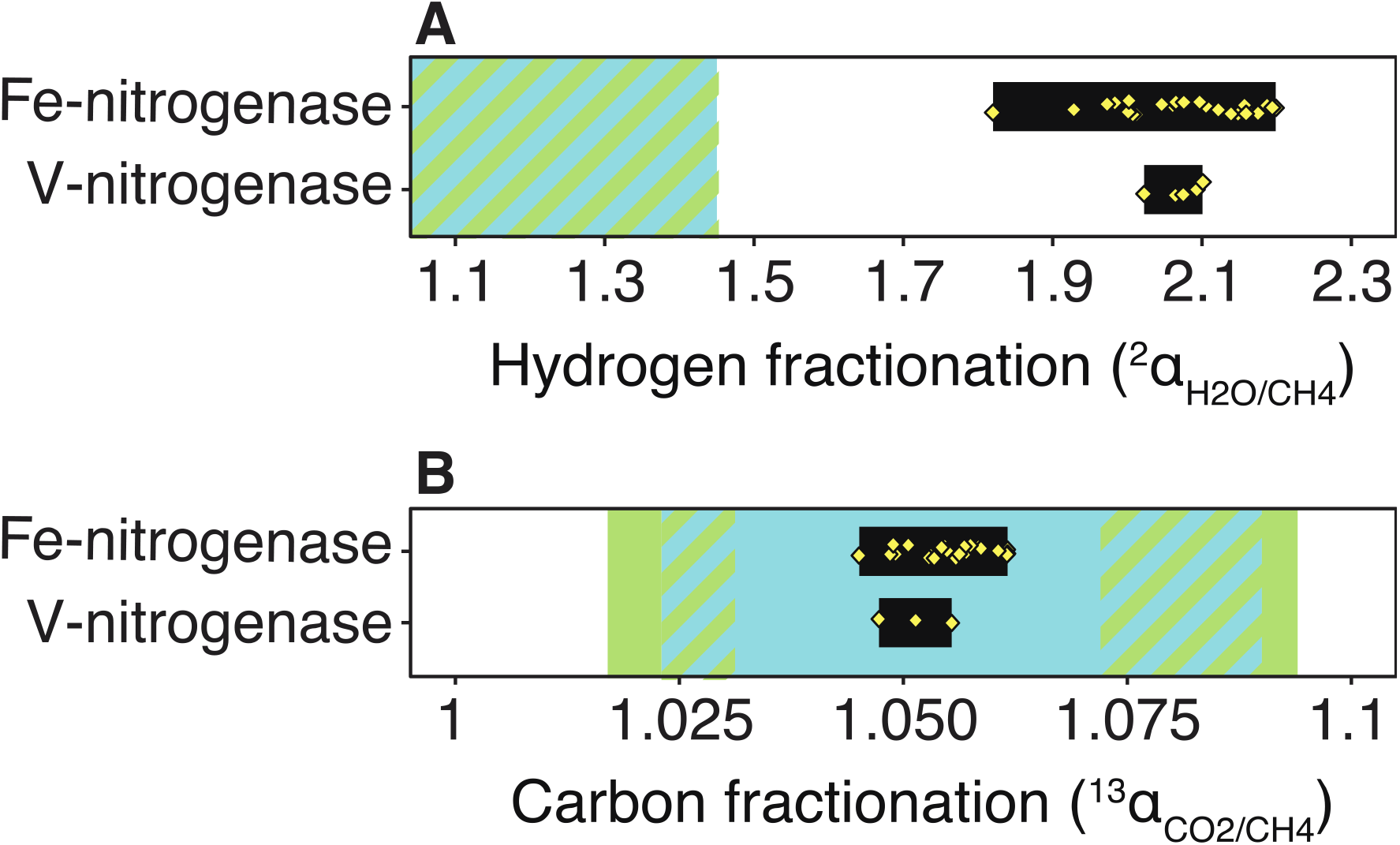
Nitrogenase-derived methane has a charachteristic hydrogen isotope fractionation. The largest hydrogen isotope fractionations observed for canonical, *mcr*-based anaerobic methanogenesis pathways, around ~ 1.45 (34, 42), are substantially smaller than the hydrogen isotope fractionation observed for nitrogenase (**A**). However, carbon isotope fractionation (**B**) by nitrogenase falls within the the range observed for hydrogenotrophic methanogenesis (blue; 1.023 ≤ ^13^α_CO2/CH4_ ≤ 1.090) and is intermediate to the range observed for methanol-(1.072 ≤ ^13^α_CO2/CH4_ ≤ 1.094) and acetate- (1.017 ≤ ^13^α_CO2/CH4_ ≤ 1.031) based fermentative methanogenesis (green; 8). Green-blue hatched areas represent ranges of overlap between the fractionation obuserved for fermentative and hydrogenotrophic methanogenesis. Individual datapoints from this study are shown as yellow diamonds (♢).

We observed only small changes in nitrogenase fractionation across a large range of temperatures, cell densities, and carbon substrates (<0.02 for ^13^α_CO2/CH4_ and < 0.38 for ^2^α_H2O/CH4_; Figs. 4, 5) relative to the variability observed for other methane production pathways. Fractionation increased by ~ 0.012 as temperature decreased from 30 to 14°C for ^13^α_CO2/CH4_ (*p* = 10^-5^) and by ~ 0.160 for ^2^α_H2O/C_H_4_ (*p* = 0.03). In contrast, the form of growth substrate (succinate or acetate) did not alter ^2^α_H2O/CH4_ (*p* = 0.96) and only had a small impact of ~0.005 on ^13^α_CO2/CH4_ (*p* = 0.006). This is compatible with the recent observation that electron availability has only a minor impact on CH_4_ production by a mutant Mo-nitrogenase isoform (44). Acidification of the growth medium by ~0.5 pH units also did not alter fractionation, though we note that there was only one biological replicate for the acidified treatment (Table 1). Despite order of magnitude differences in the rate of methane production by V- and Fe-only nitrogenase (Fig. 1), they have indistinguishable fractionation factors associated with methane production (*p* = 0.4 for ^13^α_CO2/CH4_ and 0.9 for ^2^α_H2O/C_H_4_; Table 1). This suggests there is no rate effect on fractionation and that the V- and Fe-only nitrogenases share a common mechanism for CO_2_ reduction to methane.

**Table 1.**
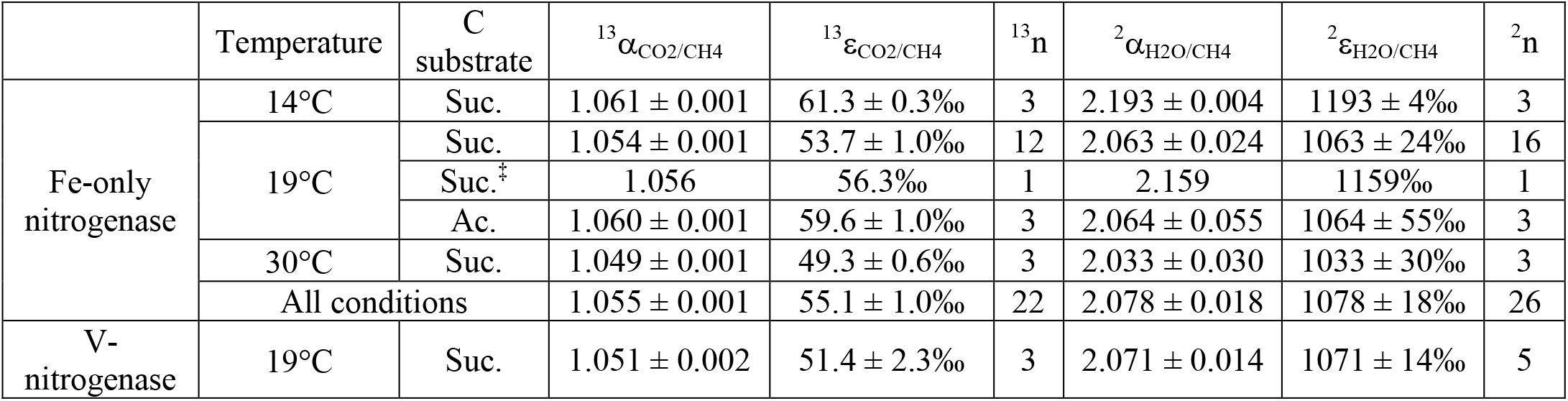
Carbon and hydrogen stable isotope fractionation associated with methane production by V- and Fe-only nitrogenase. Compared to other methane producing pathways, the range of fractionation observed over a 15°C temperature range, two different nitrogenase isoforms, and different organic carbon substrates (succinate, suc.; acetate, ac.; suc.^‡^ = acidified) is relatively small. The table shows the mean ± SE (n). Individual datapoints, including product and substrate isotopic compositions, are shown in the S.I. Table.

The greatest source of variability in fractionation (^13^α_CO2/CH4_ ~ 0.01; ^2^α_H2O/CH4_ ~ 0.25 range) appears to be due to cell density, growth phase (Figs. 5C, G), or substrate (CO_2_) concentration (Figs. 5D, H). These variables are strongly correlated due to dissolved inorganic carbon (DIC) production throughout growth (Fig. 5I) and cannot be disentangled with the current dataset.

**Fig. 5.**
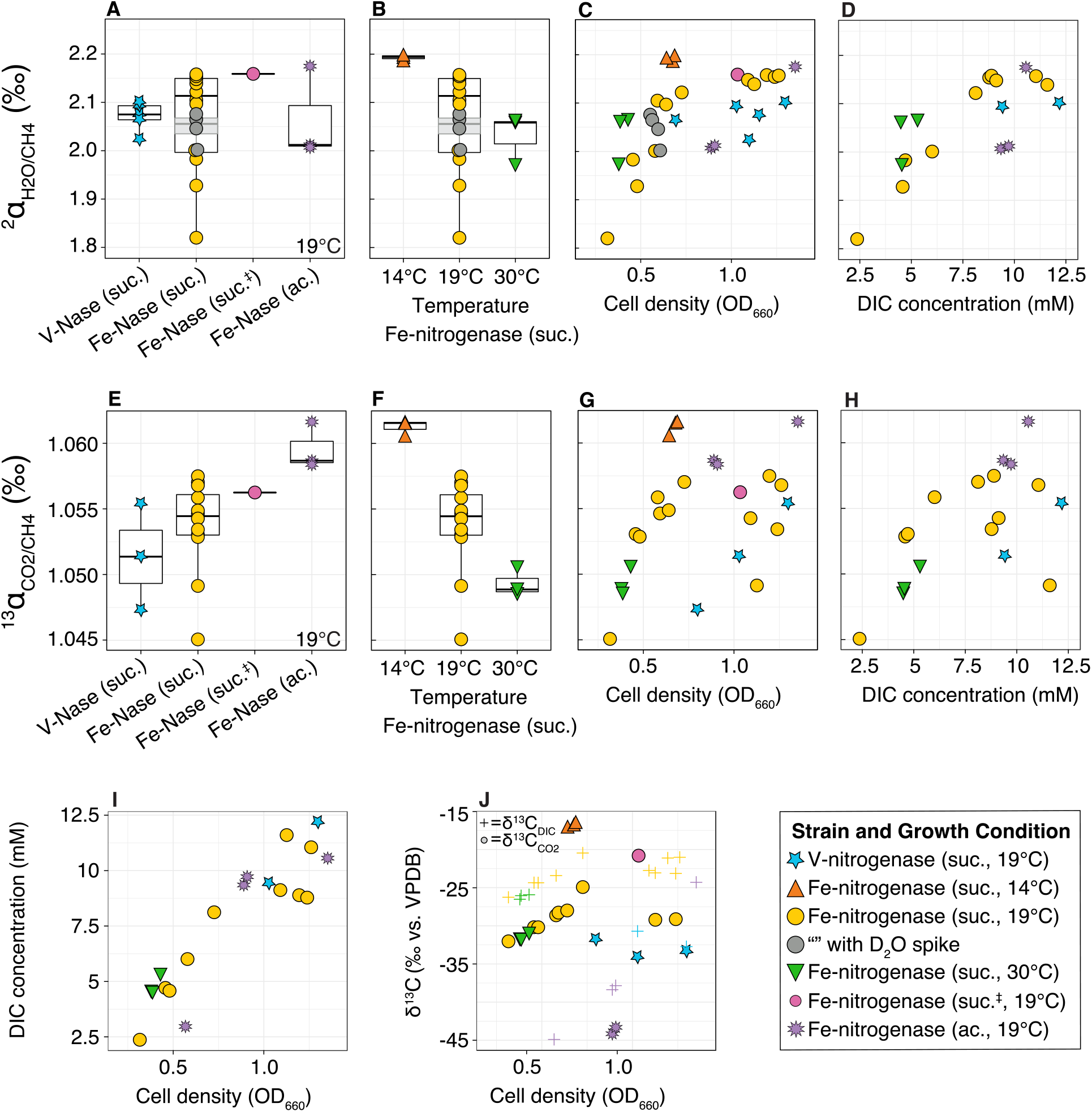
Carbon and hydrogen isotope fractionations associated with methane production by nitrogenase under different growth conditions. Hydrogen and carbon isotope fractionations increase at low temperatures (**B**, **F**), higher cell densities (**C**, **G**) and higher substrate concentrations (**D**, **H**). Hydrogen isotope fractionation is comparable during growth on different organic carbon substrates (succinate, suc.; acetate, ac.) and when the growth medium is acidified (suc.^‡^; **A**). Carbon isotope fractionation is slightly higher during growth on acetate than on succinate (**E**) although this could also be related to differences in the cell density at harvest (**G**). The dissolved inorganic carbon (DIC) concentration (**I**) and inorganic carbon isotopic composition (**J**) increase throughout exponential growth, suggesting that substrate concentration could be influencing the observed effect of cell density (OD_660_) on fractionation. The boxplots in panels **A**, **B**, **E** and **F** show the median (center line), first and third quartiles (outer lines), and values within 1.5 times the inner quartile range (whiskers). For panel **J**, note that, for some samples, only δ C_CO2_ or δ C_DIC_ were measured, such that C_CO2_ or δ C_DIC_ datapoints are not necessarily paired.

Future experiments could manipulate the DIC concentration to test the mechanistic basis for this effect. Notably, a similar cell density or growth phase effect has been previously observed for anaerobic methanogenesis, where it was tentatively attributed to changes in temperature, catabolic rate (42) or carbon assimilation during logarithmic growth (45).

The methane isotopic composition at harvest integrates the isotopic composition of methane produced throughout growth. Therefore, the fractionation measured at stationary phase is altered by the change observed in substrate CO_2_ isotopic composition during exponential phase (Fig. 5J). However, using the observed shift in medium CO_2_ isotopic composition to estimate the effect on the fractionation measured at stationary phase, we find that the change in substrate isotopic composition could account for at most half (~0.005) of the total (~0.01) shift observed in ^13^α_CO2/CH4_ with cell density (see S.I.). We note that it is possible that the isotopic composition of intracellular CO_2_ is somewhat different from the bulk composition due to localized production, consumption, and depletion, given the competing reactions of CO_2_ production during organic substrate assimilation and re-fixation by Rubisco during photoheterotrophic growth of *R. palustris* (41, 46, 47).

We observed changes in fractionation correlated with temperature, growth phase and dissolved inorganic carbon (DIC) concentration but not with organic carbon substrate or total methane production rate. Though the variability in fractionation during methane production by nitrogenase is interesting from a mechanistic perspective, the range of measured hydrogen isotope fractionation does not overlap with, and is readily distinguishable from, the range observed for other methane production pathways (Fig. 4). This is consistent with the observation that N_2_ and acetylene (C_2_H_2_) fractionations by a single nitrogenase isoform are also remarkably constant across different organisms, metabolisms and environmental conditions (26, 33).

### Hydrogen concentration does not influence methane isotope fractionation by nitrogenase

Hydrogen (H_2_) is an obligatory product of nitrogen fixation and, in our experiments, is generated simultaneously with the production of methane from carbon dioxide (48, 49). We explored whether its buildup could affect methane isotope fractionation by nitrogenase, as has been suggested for *mcr-based* methanogenesis (2, 42, 50–58). Two lines of evidence show that the presence of H_2_ does not alter the isotopic composition of methane produced by nitrogenase. Firstly, for Fe-only nitrogenase cultures (grown on succinate at 19°C in serum vials), the hydrogen isotope fractionations were indistinguishable in cultures in which the headspace contained 2-3% H_2_ at inoculation (^2^α_H2O/CH4_ = 2.068 ± 0.033, n = 3) and in cultures that were flushed with 100% N_2_ prior to inoculation (^2^α_H2O/CH4_ = 2.046 ± 0.016, n = 4, *p* = 0.57; S.I. Table). These data show that exogenous H_2_ did not influence the isotopic composition of the product methane. This result is expected given that the strains used in our experiments lack a functional uptake hydrogenase (59) and that nitrogenase itself does not catalyze isotope exchange between water and H_2_ (60). (This is a significant distinction from the hydrogenation of D_2_, forming HD, which nitrogenase ***can*** catalyze in the presence of N_2_). We note that abiotic hydrogen isotopic equilibration between H_2_-H_2_O, CH_4_-H_2_ and CH_4_-H_2_O is likely too slow to be important at the timescales (~weeks) and temperatures (≤ 30°C) of relevance to our experiments (36, 61–63). This finding is consistent with other reports that the source of protons for CO reduction by nitrogenase is water, not hydrogen gas (16).

The second line of evidence demonstrating that the hydrogen concentration does not influence nitrogenase methane isotope fractionation is based on comparing the fractionations observed in different growth containers and for the different strains. For a given growth container and strain, cell density and hydrogen concentration are correlated (Fig. 6A; also see S.I. Discussion). However, their respective effects on fractionation can be disentangled by comparing data from the Balch tubes (10 mL medium: 17 mL headspace) and serum vials (180 mL medium: 60 mL headspace). As seen in Figure 6, hydrogen and carbon isotope fractionations in cultures with 10 to 20% H_2_ in their headspace at harvest overlap with those of cultures with 20 to 50% H_2_ in their headspace at harvest (*p* > 0.5; Figs. 6C, E). We conclude that fractionation during methane production by nitrogenase is not sensitive to hydrogen concentration over the large range (10 to 50%) tested here. This is compatible with findings that CO_2_ reduction by Mo-nitrogenase is not competitively inhibited by H_2_ and does not proceed through the same reversible *re* (reductive elimination of H_2_) step as N_2_ reduction (64). The lack of hydrogen partial pressure dependency on fractionation contrasts with some modes of *mcr*-based methanogenesis.

**Fig. 6.**
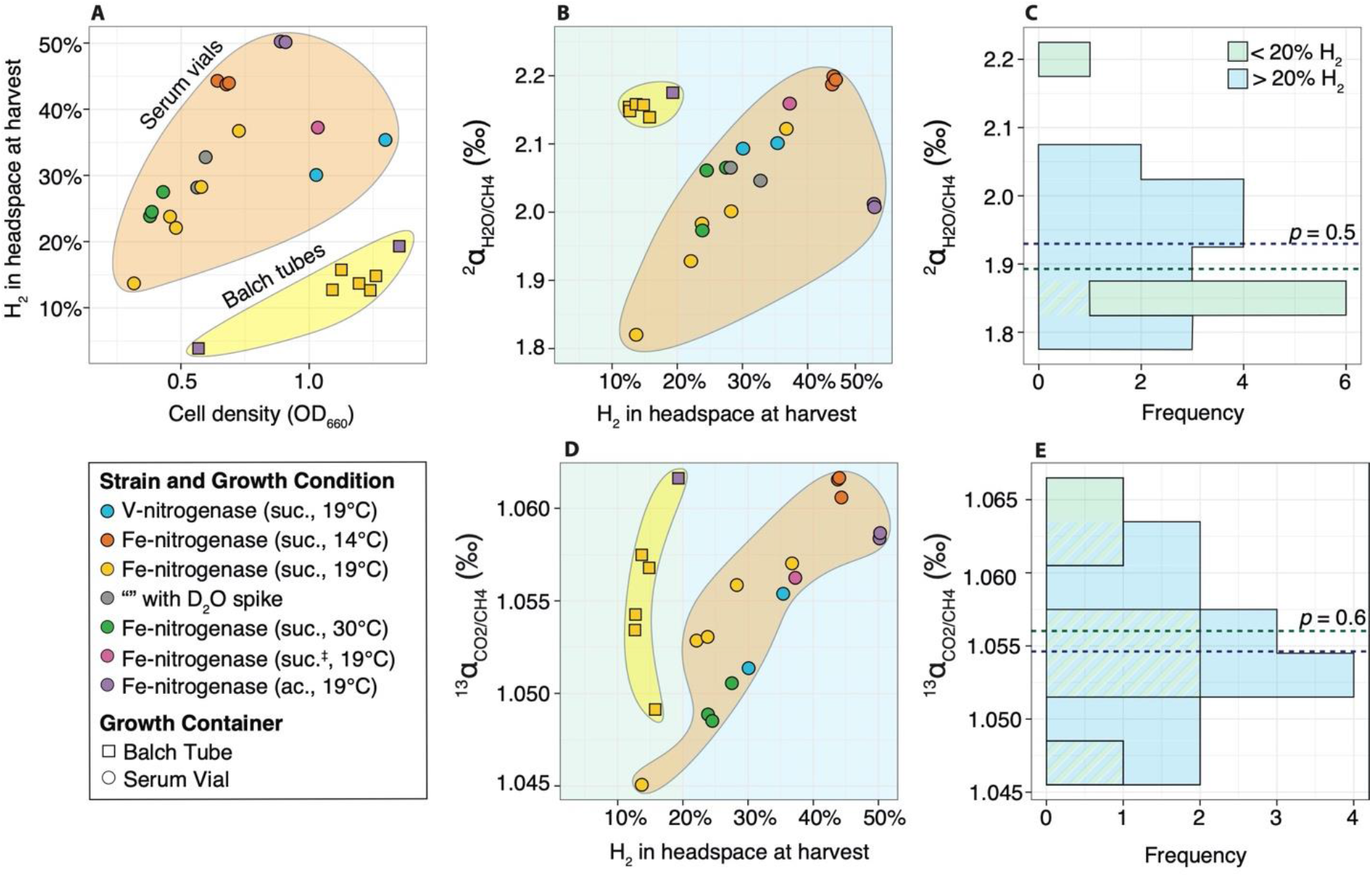
Hydrogen concentration does not alter the isotopic composition of nitrogenase derived methane. Strains produce hydrogen (H_2_) proportional to growth. Correspondingly, the cultures grown in balch tubes, which had higher headspace to volume ratios, accumulated lower concentrations of hydrogen (**A**). Comparison of hydrogen (**B**) and carbon (**D**) isotope fractionations between cultures grown in balch tubes and serum vials shows that hydrogen concentration is not responsible for the variability in fractionation between samples: the methane from cultures harvested at high cell densities in serum vials had a similar range in isotope fractionation as the methane from cultures grown in balch tubes despite >2-fold differences in headspace hydrogen concentrations. This is also apparent in histograms **C** and **E** which show that the distribution of isotope fractionation is the same (*p* > 0.5) for cultures whose headspace hydrogen concentration at harvest was between 10 and 20% (green) or 20 and 50% H_2_ (blue).

### Mechanistic implications for nitrogenase

Determining whether isotope effects are due to equilibrium or kinetic fractionation and under what conditions they are fully expressed can help elucidate the mechanism, intermediates, and reversibility of a reaction. At 20°C, the equilibrium hydrogen isotope fractionation predicted between methane and water is only ^2^α_H2O/_cH_4_ ~ 1.019 (65). This is much smaller than the fractionation observed for nitrogenase, suggesting that kinetic, rather than equilibrium, isotope effects are responsible for the large hydrogen isotope fractionation observed here. This conclusion is consistent with the finding that fractionation of CO_2_ reduction by nitrogenase is larger at colder temperatures (Fig. 5B, F), which is generally incompatible with an equilibrium isotope effect (66). These results lead us to attribute the fractionation observed here to a kinetic isotope effect (KIE) in which CH_4_ methane production by V- and Fe-only nitrogenase is roughly twice as fast as CH_3_D methane production (1.820 ≤ ^2^α_H2O/CH4_ = KIE ≤ 2.199). We suggest this new value can help yield insight into the mechanism of CO_2_ reduction by nitrogenase.

The mechanism of CO_2_ reduction by nitrogenase is a subject of much study because of its potential industrial application as a renewable fuel source (65, 67, 68 and references therein). The observation that the hydrogen KIE during methane production is ~2 represents a new experimental constraint for these studies. Previously, the KIE for H_2_ production in the absence of N_2_ (i.e. E4 to E2 state, where E2 is an intermediate state in the sequential reduction of the active site to prepare for N_2_ binding at E4) by the Mo-nitrogenase was used as a tool to determine the mechanism of H_2_ loss during activation of the cofactor, a catalytically inefficient reaction that competes with N_2_ reduction (69). Khadka and colleagues (69) demonstrated, experimentally and computationally, that the KIE of ~2.7 is due to preference for ^1^H during protonation of the bridging Fe-hydrides by highly acidic, protonated cofactor thiols. The KIE of ~2 observed here is lower than the KIE measured for H_2_ production. This hints that (1) the preference for ^1^H might be somewhat lower for V- and Fe-only nitrogenase compared to the Mo-nitrogenase (e.g., (70–72) and references therein for examples of the effect that the cofactor and amino acid environment have on protonation and substrate selectivity). Another possibility (2) is that, because the mechanism of CO_2_ reduction by nitrogenase likely involves the migratory insertion of cofactor bound CO_2_ into the Fe-hydride bond (64), the preference for ^1^H is lower for the bridging Fe-hydrides, which do not exchange with solvent at the timescales relevant to the reaction, compared to protonated thiols, which do (69). It is also possible that proton tunneling, which is generally thought to have a very large kinetic isotope effect (but also see 73, 74) and has been proposed to occur in nitrogenase (75) could be contributing to the observed KIE, though we note that the temperature effect observed here is opposite of the predicted effect for tunneling (76, 77). Computational models can distinguish the rates of hydrogenation based on ^1^H and ^2^H and might be able to shed light on whether currently proposed, multi-step mechanisms of hydrogenation by nitrogenase (78–81) are compatible with the measured KIE of ~2. The clumped isotopic composition of methane produced by nitrogenase could also provide additional constraints.

### Environmental Relevance

The carbon and hydrogen isotopes of methane are critical constraints for the attribution of emissions of this potent greenhouse gas to its sources (82). Our characterization of nitrogenase’s biosignature helps refine the space of possible source δ^13^C and δ^2^H values. The characteristic δ^2^H signature of alternative nitrogenases distinguishes them from other microbial and thermogenic methane sources (Fig. 2). At −550%₀, the δ^2^H of nitrogenase-derived methane falls well below the lowest values, around −400%₀, that have been observed for other biotic and abiotic processes (2, 38).

Given the ubiquity of CO_2_ in cells and in the environment, it is likely that some CH_4_been attributed to the canonica lproduction is occurring whenever V- and Fe-nitrogenase are active. To determine the extent to which stable isotopes can attribute methane production to alternative nitrogenase activity in environments with multiple sources, we developed a simple isotopic mixing model (Fig. 7). The model calculates the net ^2^α_H2O/CH4_ and δ^2^H of the mixed methane pool given the local water isotopic composition and the relative rates of methane production from traditional methanogenesis pathways and nitrogenase activity, assuming that all hydrogen for acetoclastic methanogenesis ultimately derive from environmental water. Hydrogen isotopic compositions as low as δ^2^HCH4 ~ −400%₀ have been attributed to the canonical hydrogenotrophic and acetoclastic methanogenesis pathways in natural samples (2, 38). Using this value as an upper bound, we suggest that measured ^2^α_H2O/CH4_ ≥ 1.65 (shown in red in Fig. 7A) would provide evidence for alternative nitrogenase activity in natural samples. Thus, the isotopic mixing model demonstrates that methane stable isotopes can identify alternative nitrogenase activity as long as the rate of methane production from nitrogenase is faster or in the same range as anaerobic, *mcr*-based methanogenesis rates (nitrogenase methane: total methane > 0.5) and that the isotopic composition of source water must be taken into account when interpreting the relative contributions of different biotic methanogenesis pathways. The model provides quantitative bounds on the use of characteristically low δ^2^H of methane produced by nitrogenase as a biosignature of alternative nitrogen fixation.

**Fig. 7.**
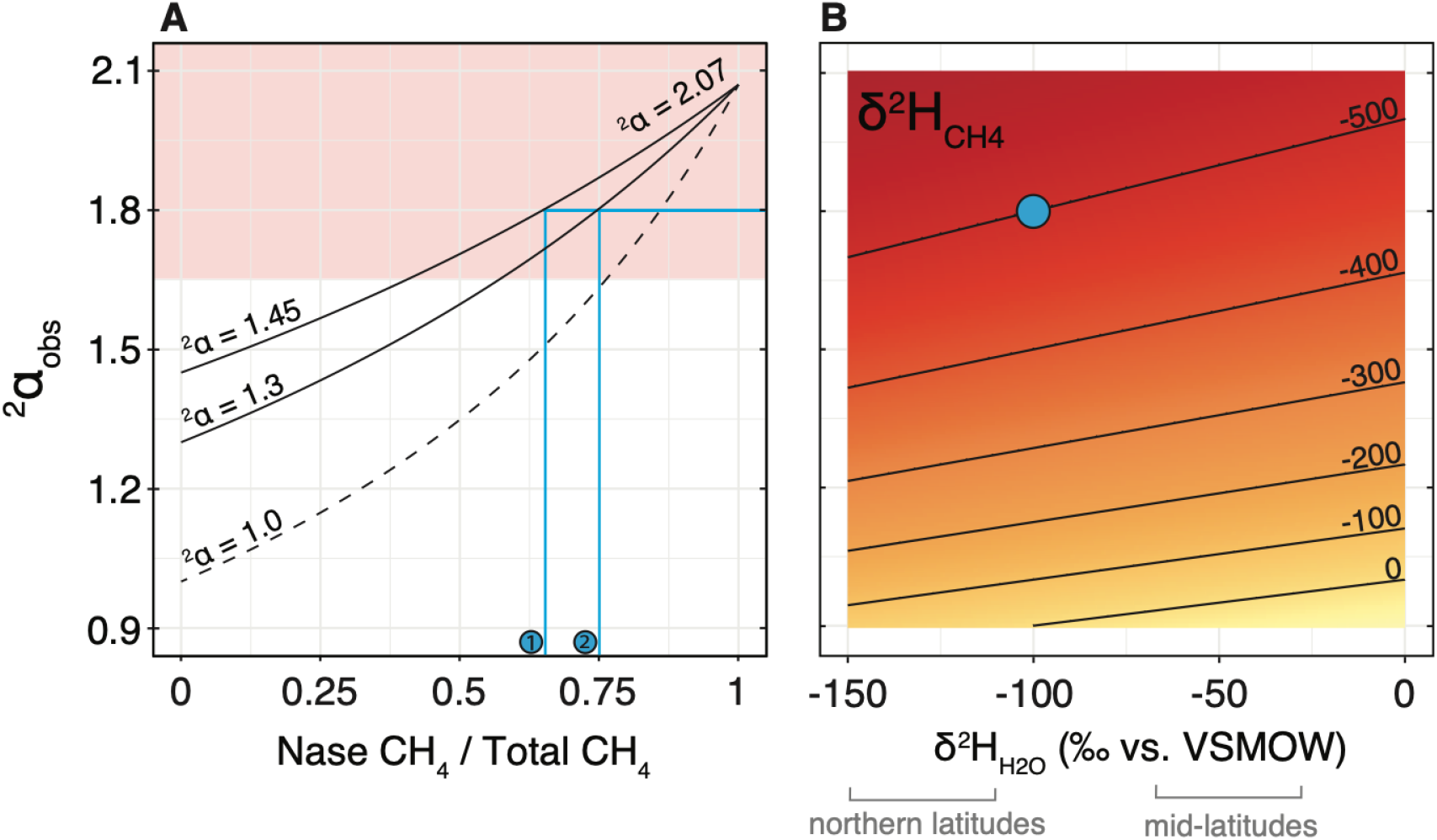
Observed fractionation (A) and methane isotopic composition (B) for multiple methane sources. Panel A shows the apparent fractionation (^2^α_H2O/CH4_) between water and methane in an environment with co-occurring production from nitrogenase (^2^α_H2O/CH4_ = 2.07; x-axis shows the relative contribution of nitrogenase to the total methane pool) and a second source with ^2^α_H2O/CH4_ = 1.0 (dashed line; no expressed fractionation), ^2^α_H2O/CH4_ = 1.3 (broadly representative of hydrogenotrophic methanogenesis (34)) or ^2^α_H2O/CH4_ = 1.45 (broadly representative of acetoclastic methanogenesis (34), though values this high have also been observed for hydrogenotrophic methanogenesis (42)). Panel B shows the calculated methane isotopic composition δ^2^HCH4 given the isotopic composition of local source water (x-axis) and the measured fractionation between water and methane (^2^α_H2O/CH4_; y-axis). For example, for a sample with ^2^α_obs_ = 1.8 (e.g. when δ^2^H_H2O_ = 100%₀, δ^2^H_CH4_ = −500%₀, shown as a blue circle in panel B) the model predicts that ≥ 70% of the methane is produced by nitrogenase, depending on whether the competing source of methane is fermentative (blue line 1, ~70%) or hydrogenotrophic (blue line 2, ~75%).

It is clear that methane production by the V- and Fe-only nitrogenases does not contribute quantitatively to methane production at the global scale (10). For instance, assuming generously that ~20% of the ~145 Tg annual terrestrial biological nitrogen fixation flux (~120 Tg year^-1^ from (83) corrected for underestimation by the acetylene reduction assay as described in (26)) is fixed by Fe-only nitrogenase, and recognizing that methane itself is a minor byproduct of dinitrogen reduction (~ 5 *x* 10^-4^ CH_4_: 1 N_2_ for Fe-only nitrogenase, data not shown), the resultant ~0.01 Tg year^-1^ is negligible compared to total methane emissions of ~560 Tg year^-1^ (84). Nonetheless, we hypothesize that it could influence the methane isotopic composition, and act as a biomarker for alternative nitrogenase activity, in nitrogen-limited environments with low methanogenesis rates and high alternative nitrogenase activity. The controls on alternative nitrogenase activity are not fully understood (e.g. Glazer *et al.,* 2015; Zhang *et al.,* 2016), although new tools (26, 33) are rapidly advancing our understanding of their distribution. It is now well established that alternative nitrogenases are favored under conditions of low Mo availability (27, 86), though their activity has been observed in some sedimentary environments that appeared to be Mo-replete as well (25, 26). Aerobic soils, cyanolichens, mosses and other biocrusts, lake and marine waters (8, 87), or sediment systems with high sulfate concentrations, where sulfate reducers generally outcompete methanogens for substrates (3, 88), are possible targets to test when alternative nitrogenases are active using methane stable isotopes (10). We note that, in the global inventory of methane isotopic data, the single lowest δ^2^H composition of −442%₀ was recorded during the fall in northern Canada (38, 89), which coincides seasonally and spatially with measurements of high alternative nitrogenase activity in boreal cyanolichens (27, 28). This presents an exciting avenue for future research aimed at constraining the importance of nitrogenase to methane production in environments with low activity of canonical methanogens, and at illuminating the mechanism(s) of nitrogenase CO_2_ reduction.

## Conclusion

The alternative V- and Fe-only nitrogenases are important enzymes in the global nitrogen cycle. The curious observation that these enzymes produce methane as a minor byproduct of nitrogen fixation led us investigate how its isotopic compostion compares to other natural methane sources. Here we show that the natural abundance deuterium to hydrogen ratio of methane derived from nitrogenase is significantly lower than methane from all other known processes, with δ^2^H as low as −550%₀. This result provides new experimental constraints on the mechanism of the nitrogenase enzyme and demonstrates that significantly depleted hydrogen stable isotopic composition constitute a passive biosignature of V- and Fe-only nitrogenase-derived methane. This isotopic fingerprint offers a means to probe the contribution of alternative nitrogen fixation and nitrogenase methane emissions on Earth and beyond.

## Materials Methods

### Bacterial cultures

*Rhodopseudomonas palustris* strains CGA766 (“V-nitrogenase strain,” genotype: *Δniufll nifD::Tn5 ΔanfA*) and CGA755 (“Fe-only nitrogenase strain,” genotype: *ΔnifH ΔvnfH*) were grown in batch cultures at 14, 19 and 30°C and ~90 μmols photons m^-2^ s^-1^ under anaerobic photoheterotrophic conditions in defined nitrogen-fixing medium with 2.5 μM Fe, 100 nM Mo, 10 μM V, Wolfe’s vitamin solution, 0.0005% yeast extract and either 10 mM succinate or 20 mM acetate (33, 40, 41). Where applicable, the δ^2^H of the growth medium was manipulated by adding 99.9% purity D_2_O (Cambridge Isotope Laboratories, Inc.) to the growth medium. Bacterial growth was monitored by optical density (OD_660_) using a Genesys 20 visible spectrophotometer (Thermo Fisher Scientific) and converted to cell density using the empirically observed relationship cells mL^-1^ = 2.29 *x* 10^9^ *x* OD_660_.

### Analytical

Methane concentrations in the culture headspaces were measured either on a Peak Performer 1 gas chromatograph with N_2_ carrier gas (Peak Laboratories) or on a GC-8A with He carrier gas (Shimadzu Instruments; column = Supelco HayeSep N; column temperature = 80°C; detector temperature = 150°C) with flame ionization detectors. Calibration curves were made by sequentially diluting 100 ppm or 1% CH_4_ standards with N_2_ in a 10 mL syringe with a luer-lock and, like the samples, loading 1 mL onto the instrument using an injection loop. Hydrogen and carbon dioxide gas concentrations were measured using gas chromatography with a thermal conductivity detector (GC-8AIT TCD, Shimadzu Instruments; column = Restek ShinCarbon ST; column temperature = 100°C; detector temperature = 150°C) with N_2_ as the carrier gas. Dissolved methane was not quantified. We note that not all variables were measured in all samples, and that the raw datapoints used for all the figures and calculations in this manuscript are available in the S.I.

### Stable Isotope Measurements

Methane samples were analyzed for δ^2^H and δ^13^C at the UC Davis Stable Isotope Facility. Depending on the methane concentration, samples were collected either in pre-evacuated 12 mL soda glass vials (Labco Limited; 839W) or diluted in He-flushed vials. Because sample methane δ^2^H was depleted relative to the lowest standard available at the UC Davis Stable Isotope Facility (−276‰), a dilution series of a single sample was measured, and the resulting linearity correction applied to all samples (calculations included in the S.I. Table). The constant hydrogen isotope fractionation observed for Fe-only nitrogenase over a >500‰ range in δ^2^H suggests that the analytical methods employed are robust (Fig. 3). Samples for δ^13^C analysis of CO_2_ were collected in the same manner as those for methane. Samples for δ^13^C of DIC were collected in He-flushed vials that contained 1 mL of concentrated HPLC grade phosphoric acid (85%; Fisher Chemical). At the UC Davis Stable Isotope Facility, the δ^2^H_CH4_, δ^13^Cc_H4_, δ^13^C_CO_2__ and δ^13^C_DIC_ samples were measured on a Delta V Plus IRMS (Thermo Scientific, Bremen, Germany) coupled to a Gas Bench II system. Water δ^2^H samples were collected by filtering growth medium (0.22μm) at the end of the experiment and storing at −20°C. For analysis, samples were thawed and 1.4 – 1.5 mL were aliquoted into 2 mL soda glass vials (Thermo Scientific, National C4010-1W with C4010-40A caps) and shipped on ice or at room temperature overnight to the UC Davis Stable Isotope Facility, where they were measured on a Laser Water Isotope Analyzer V2 (Los Gatos Research, Inc.). Biomass and substrate δ^13^C were measured in the Zhang stable isotope laboratory at Princeton as described previously (41) on a Vario ISOTOPE select (Elementar Isoprime). The standard deviation of standard material replicates were < 1% for δ^2^H_H2O_, < 2% for δ^2^H_CH4_, < 0.2% for δ^13^C_CH4_ (> 10 ppm), < 0.2% for δ^13^CCO_2_ and δ^13^C⊓Ic, and < 0.1% for δ^13^C_biomass_.

### Isotope Calculations

Hydrogen and carbon isotopes are expressed using delta notation relative to Vienna Standard Mean Ocean Water (VSMOW) and Vienna Pee Dee Belemnite (VPDB), respectively. Apparent CO_2_-CH_4_ and water-CH_4_ isotope fractionation factors were calculated as substrate over product using the equations:

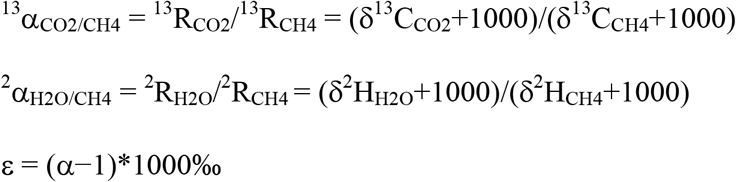

In this manuscript, errors represent the standard error of multiple biological replicates.

### Isotope Mixing Model

To determine under what conditions the methane isotopic composition can be used as a biosignature for alternative nitrogenase activity, we developed a mixing model that calculates the fractionation and isotopic composition of methane produced by multiple sources (Fig. 7). We used the following parameters: ^2^α_Nase_ = 2.07; δ^2^H_H2O_ = −40%₀ vs. VSMOW as representative of the mid-latitudes and −150%₀ vs. SVMOW as representative of northern latitudes; and *k* = methane produced by nitrogenase: total methane produced by nitrogenase and *mcr*-based anaerobic methanogenesis. For fermentative methanogenesis, the model assumes that all protons ultimately derive from local water. The observed fractionation and isotopic composition were calculated using the equations:

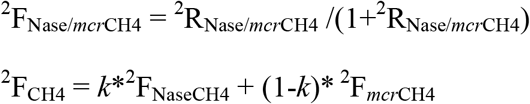

### Data Availability

Individual datapoints are available in the S.I. Table. In addition to the S.I. Table, these data will be uploaded in FigShare prior to publication.

## Acknowledgements

We thank Richard Doucett and Elvira Delgado of the UC Davis Stable Isotope Facility for the isotopic analysis of the water, methane, CO_2_ and DIC samples in this project, and Lina Taenzer, Rachel Harris and Barbara Sherwood Lollar for useful discussions. Ashley Maloney and Emma Bertran provided valuable feedback on drafts of this paper. Funding for this project was provided by the National Science Foundation and National Aeronautics and Space Administration (NSF Award# EAR1631814 and NASA Award# 80NSSC17K0667 to XZ), an NSF Graduate Research Fellowship to KEL, the Princeton Environmental Institute through the Walbridge Fund, the Simons Foundation division of Life Sciences (XZ, WDL), and the Dartmouth College Vice Provost for Research (WDL).

